# A Panel of Diverse *Klebsiella pneumoniae* Clinical Isolates for Research and Development

**DOI:** 10.1101/2022.08.17.504361

**Authors:** Melissa J. Martin, William Stribling, Ana C. Ong, Rosslyn Maybank, Yoon I. Kwak, Joshua A. Rosado-Mendez, Lan N Preston, Katharine F. Lane, Michael Julius, Anthony R. Jones, Mary Hinkle, Paige E. Waterman, Emil P. Lesho, Francois Lebreton, Jason W. Bennett, Patrick T. McGann

## Abstract

*Klebsiella pneumoniae* are a leading cause of healthcare associated infections worldwide. In particular, strains expressing extended-spectrum β-lactamases (ESBLs) and carbapenemases pose serious treatment challenges, leading the World Health Organization (WHO) to designate ESBL and carbapenem-resistant Enterobacteriaceae (CRE) as “critical” threats to human health. Research efforts to combat these pathogens can be supported by accessibility to diverse and clinically relevant isolates for testing novel therapeutics. Here, we describe a panel of 100 diverse *K. pneumoniae* isolates publicly available to assist the research community in this endeavor.

Whole-genome sequencing (WGS) was performed on 3,878 *K. pneumoniae* clinical isolates housed at the Multidrug-Resistant Organism Repository and Surveillance Network. The isolates were cultured from 63 facilities in 19 countries between 2001 and 2020. Core-genome multilocus sequence typing and high-resolution single nucleotide polymorphism based phylogenetic analyses captured the genetic diversity of the collection and were used to select the final panel of 100 isolates. In addition to known multi-drug resistant (MDR) pandemic lineages, the final panel includes hypervirulent lineages and isolates with specific and diverse resistance genes and virulence biomarkers. A broad range of antibiotic susceptibilities ranging from pan-sensitive to extensively drug resistant isolates are described. The panel collection, all associated metadata and genome sequences, are available at no additional cost and will be an important for the research community and for the design and development of novel antimicrobial agents and diagnostics against this important pathogen.

**Importance:** *Klebsiella pneumoniae* is a major cause of healthcare-associated infections that are increasingly difficult to treat due to the emergence of multi-drug resistant strains. In particular, strains expressing extended-spectrum β-lactamases and carbapenemases have attained global notoriety, with the World Health Organization listing these strains as a “critical-priority” for the development of new therapeutics. Access to a diverse collection of strains for testing is critical for this endeavor, but few resources currently exist. Similarly, pivotal research of the genetic determinants underlying the pathogenesis of hypervirulent lineages is hampered by the lack of standardized, comparator strains. Herein we describe a panel of 100 diverse *K. pneumoniae* constructed to maximize genetic and phenotypic diversity from a repository of over 3,800 clinical isolates collected over 19 years. The panel, and all associated metadata and genome sequences, is provided at no cost and will greatly assist efforts by academic, government, and industry research groups.

## Introduction

*Klebsiella pneumoniae* are a leading cause of nosocomial infections resulting in pneumonia, bacteremia, surgical site, and urinary tract infections (1). A member of the problematic ESKAPE (*Enterococcus faecium, Staphylococcus aureus, Klebsiella pneumoniae, Acinetobacter baumannii, Pseudomonas aeruginosa, Enterobacter)* group of pathogens (2), “classical” *K. pneumoniae* (cKp) are associated with prolonged outbreaks, increased disease burden, and high mortality rates (3, 4). The prevalence of cKp infections has steadily increased since 2005, primarily driven by strains acquiring extended-spectrum β-lactamases (ESBLs) and carbapenemases conferring resistance to 3^rd^ generation cephalosporins and carbapenem antibiotics (5, 6). These multidrug resistant (MDR)-cKp clones are a threat to the medical community as antibiotic treatment options are limited and non-susceptibility to all antibiotics has been reported (7). In alignment, the World Health Organization (WHO) ranks *K. pneumoniae* among the critical priority list for the development of therapeutics (8).

In parallel to hospital-acquired MDR-cKp, severe community-acquired infections caused by so called “hypervirulent” *K. pneumoniae* (hvKp) lineages have also emerged (9). These invasive strains are generally susceptible to antibiotics and generally occur in healthy hosts causing meningitis, liver abscesses, endophthalmitis, and soft tissue infections (9). hvKp strains are associated with the acquisition of large virulence plasmids and/or mobile elements encoding virulence determinants such as siderophores [e.g aerobactin (*iuc*), salmochelin (*iro*), yersiniabactin (*ybt*)], metabolite transporter *peg-344*, genotoxic polyketide colibactin (*clb*), and regulators of mucoviscosity and capsular polysaccharide (*rmpA* and *rmpA2*) (10, 11). While there are distinct clinical and genetic differences between the two main pathotypes of *K. pneumoniae*, there has been a concerning emergence of convergent lineages that carry both MDR and virulence determinants (12–14). This confluence of MDR-cKp and hvKp has provided additional impetus to develop novel antibiotics and therapeutics (15).

The *K. pneumoniae* population is diverse consisting of over 250 clonal phylogenetic lineages and an estimated accessory genome of >100,000 protein coding sequences (6, 16). Despite hundreds of clones that can cause infections, a few “high-risk”, globally disseminated, MDR-cKp lineages (e.g. ST-11, ST-14, ST-101, ST-147, ST-258, ST-307) contribute to the majority of infections (6). For example, the dissemination of KPC-type carbapenemases is largely attributed to the well-studied, clonal ST-258 lineage, which is now endemic in many countries, including the United States (17–19). More recently, carbapenem resistant ST-307 and ST-147 clonal lineages carrying various carbapenemases (NDMs, OXA-48-like, and KPC) have emerged and are circulating in countries such as the United States (19), Germany (20), and in Italy (21). In contrast, unrelated hvKp lineages are mainly described from the Asian Pacific Rim countries and are predominately ST-23, ST-86, ST-65, ST-380, and ST-66 lineages (6, 9). These hvKp strains are associated with very few capsular polysaccharide types K1, K2, and/or K5, in contrast to the substantial diversity of K-loci found in cKp strains (22). The significant genomic diversity and constantly changing epidemiology highlights the importance of using the *K. pneumoniae* population structure for identifying diverse isolates when developing effective targets for treatments and diagnostics against problematic MDR-cKP, hvKp, and emerging clones.

In this report, we utilized the large repository of 3,878 clinical *K. pneumoniae* maintained by the Multidrug-Resistant Organism Repository and Surveillance network (MRSN) (23) and collected globally between 2001 and 2020. Comparable to our previous work (24, 25) we constructed a reference panel of 100 *K. pneumoniae* clinical isolates that captures the extensive genetic diversity of this species, as well as variable antibiotic resistance gene content and virulence gene content along with a wide range of antimicrobial susceptibility profiles. This panel is available to the research community at no extra cost to aid in the design and development of novel therapeutics and diagnostics for this critical pathogen.

## Results

### Global *K. pneumoniae* population structure and collection diversity

3,878 *K. pneumoniae* clinical isolates were collected over a 19-year period (2001 to 2020) from across the U.S. and globally in collaboration with the U.S. Department of Defense’s Global Emerging Infections Surveillance (GEIS) branch. After removal of serial isolates from the same patients, 3,123 primary isolates from 2,760 patients were analyzed by core-genome multilocus sequence typing (cgMLST) to generate a minimum spanning tree revealing the genomic diversity of the population (**Fig. 1A**). The isolates were recovered from 63 healthcare facilities across 6 continents including North America (63%), Asia (17.6%), Europe (8.9%), South America (5.0%), Africa (4.7%), and Oceania (0.4%). The majority were cultured from urine (46%), followed by respiratory (11%), perianal surveillance swabs (10%), wound (9%), blood (9%), and body fluid (2%) cultures. *In silico* MLST using the scheme designed by Diancourt *et al*. (26) identified 480 ST’s with 260 (54%) found in isolate(s) from a single patient. Despite the large number of ST’s, 34% of the isolate collection is represented by 6 globally problematic clones: ST-15 (7.8%), ST-147 (5.9%), ST-258 (5.8%), ST-307 (5.3%), ST-14 (4.8%), and ST-16 (4.6%) (6). Clonal lineages were associated with lower allelic diversity (e.g. ST-258 maximum of 87 allelic differences) however, extensive diversity was observed within other lineages (e.g. 1,312 allelic differences within ST-37) **(Fig. 1A**).

**Figure 1.**
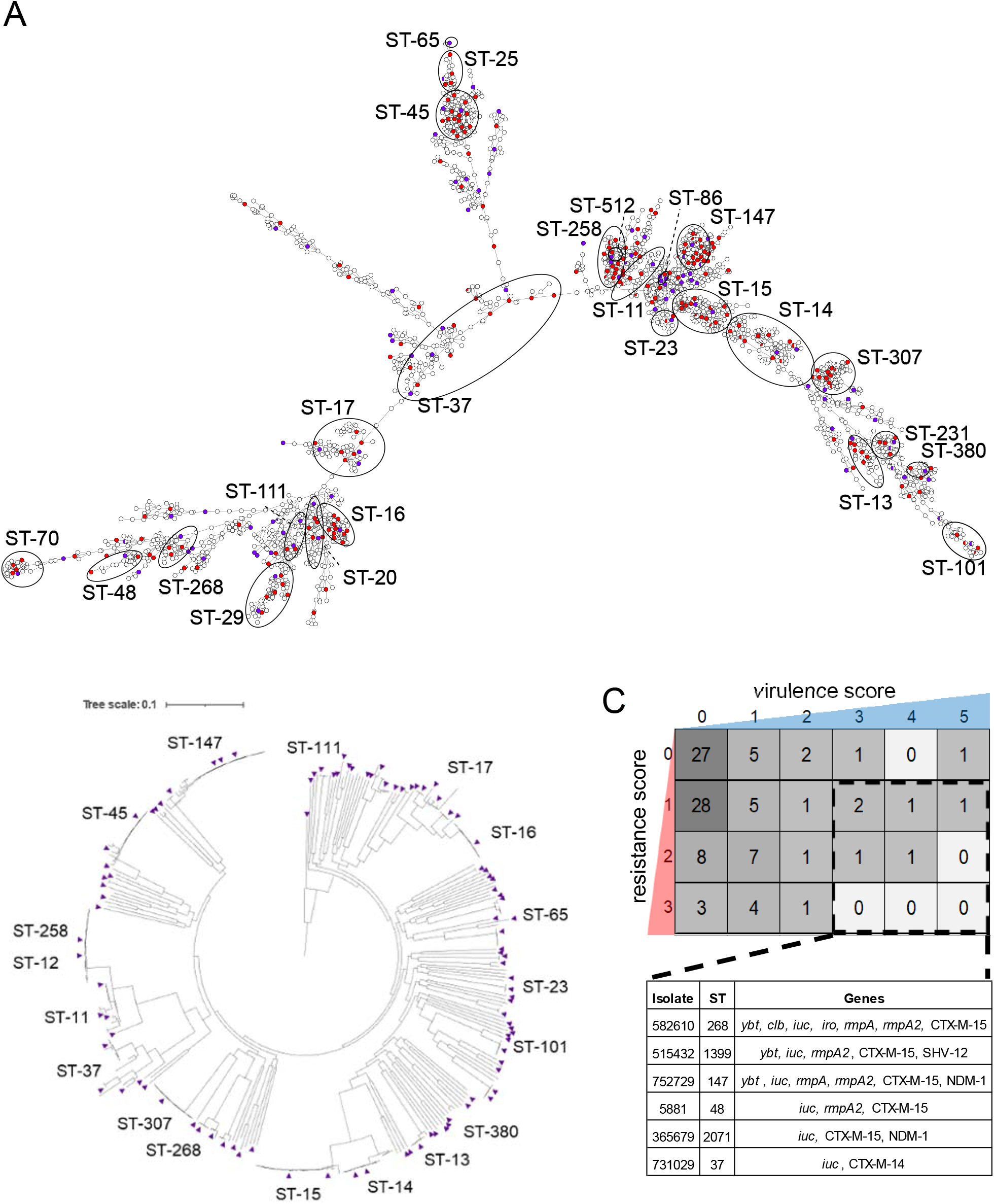
Genomic diversity of *K. pneumoniae* in the MRSN collection. A) cgMLST minimum spanning tree of the 3,123 *K. pneumoniae* genomes. Isolates with an identical MLST profile are represented within a single circle. Initial subset of isolates selected are indicated by filled red circles (*n* = 346) and the final panel isolates are indicated by filled purple circles (*n* = 100). B) Core genome SNP phylogenetic tree of 346 *K. pneumoniae* isolates initially selected to represent the breadth of *K. pneumoniae* diversity. The final 100 isolates selected for the panel are indicated in purple triangles. C) Heatmap indicating the combination of virulence/resistance scores for all panel isolates. *In silico* prediction using the Kleborate typing tool and visual inspired from Lam and coauthors (54). The number of isolates with a specific score are indicated in the boxes. Convergent isolates are indicated by the dashed black box and listed in the table below. All convergent isolates are carrying the *iuc* loci and an ESBL and/or carbapenemase gene.

### Selection of a nonredundant, genetically diverse panel of *K. pneumoniae*

Based on the cgMLST analysis, an initial subset of 346 isolates (11%) was selected to represent the maximum genetic diversity of the collection and to minimize clonal redundancy (**Fig. 1A, red dots**). This subset, encompassing 143 STs, was further compared using a maximum likelihood single nucleotide polymorphism (SNP)-based phylogenetic tree (**Fig. 1B**). In an effort to provide a pragmatic panel, 100 isolates were selected from the subset and analyzed by core genome SNP-based phylogeny (**Fig. 2 and Table S1**). This final panel of 100 isolates encompassed 94 STs, including 6 novel ST’s, and retained substantial diversity in gene content. The core genome encompassed 3,034 genes with the pangenome consisting of 21,419 genes (**Fig. S1**). Similar to previous studies (27, 28), the most prevalent predicted O antigen types, involved in the composition of cell surface lipopolysaccharide, were O1, O2, and O3, which were found in 81% of the panel isolates, followed by types O4 (10%), O5 (5%), and unknown (4%) (**Table S1**). The panel also contains 54 distinct capsular polysaccharide types, with K2 type being the most prevalent (*n* = 7).

**Figure 2.**
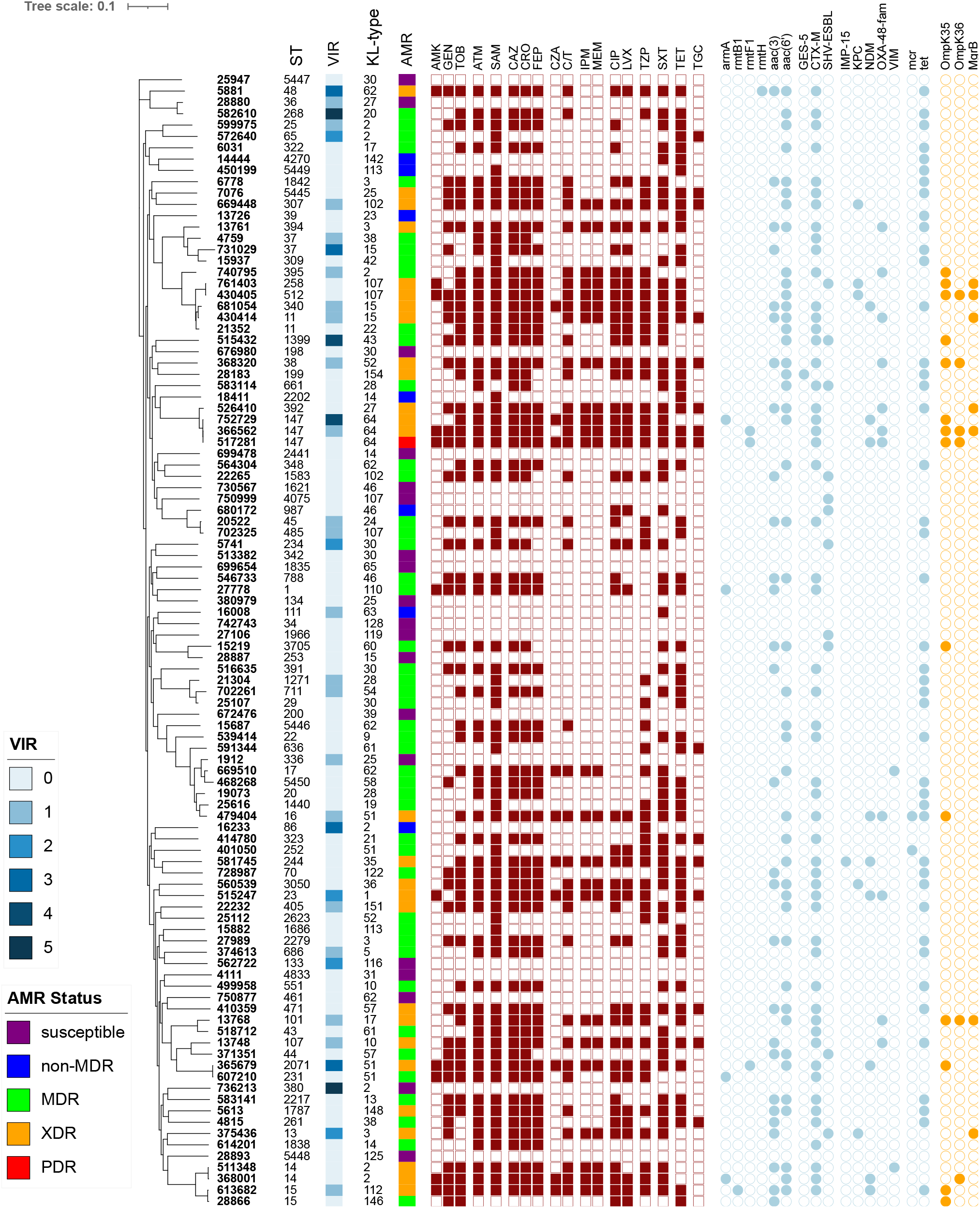
Characteristics of the *K. pneumoniae* diversity panel. Core genome SNP-based phylogenetic tree of the 100 genomes in the final diversity panel. Sequence-type (ST), virulence score (see legend), capsule polysaccharide locus, KL-type, and AMR status (see legend and text for additional details) are indicated in the columns. The assigned antimicrobial resistance phenotype for each antibiotic tested is indicated by the maroon squares-a result of non-susceptible (filled) or susceptible (open). The light blue circles indicate the presence of a known antimicrobial resistance gene, and the orange circles indicate the presence of a known mutation/truncation. AMK, amikacin; GEN, gentamicin; TOB, tobramycin; ATM, aztreonam; SAM, ampicillin/sulbactam; CAZ, ceftazidime; CRO, ceftriaxone; FEP, cefepime; CZA, ceftazidime/avibactam; C/T ceftolozane/tazobactam; IPM, imipenem; MEM meropenem; CIP, ciprofloxacin, LVX, levofloxacin; TZP, piperacillin/tazobactam; SXT, Sulfamethoxazole-Trimethoprim; TET, Tetracycline; TGC, Tigecycline.

### Distinct virulence gene content in the *K. pneumoniae* panel isolates

Acquired *K. pneumoniae* virulence loci associated with the hvKp pathotype were characterized in the panel isolates. Thirty isolates in the final panel carried the *ybt* siderophore gene cluster found on chromosomally inserted integrative conjugative elements (ICE*Kp*). The ICE*Kp*3 lineage (encoding *ybt*9 sequence type) was the most prevalent and found in 8 isolates from ST-11, ST-15, ST-16, ST-101, ST-147, ST-307, ST-340, and ST-1271 (**Table S1**). Seven isolates carried *clb*, encoding the genotoxic colibactin, in conjunction with *ybt* (lineage 1, 12, and/or 17) that were associated with the ICE*Kp*10 lineage, as previously described (29). The *iro* gene cluster encoding salmochelin synthesis and the regulators for hypermucoidy and capsule expression, *rmpA* and/or *rmpA2*, were identified in 3 isolates. Notably, 8 isolates harbored the aerobactin-encoding *iuc* genes, including 2 known hvKp lineages (ST-380 and ST-86) with the predicted serum resistant K2 capsular serotype (22). Further, 6 isolates carried the *iuc* loci in addition to ESBL genes (*bla*_CTX-M-14_ or *bla*_CTX-M-15_) (**Fig. 1C)**. Alarmingly, 2 of these genotypic convergent isolates also carried the *bla*_NDM-1_ carbapenemase including the recently characterized epidemic ST-147 isolate, MRSN 752729, from a nosocomial outbreak in Italy (12).

### AMR gene content and antimicrobial susceptibilities of the final panel

64 distinct antibiotic susceptibility profiles were observed in the final 100 isolate panel (**Fig. 2 and Table S1**). Using the susceptibility criteria developed by Magiorakos *et al*. (30), 1 isolate was pan drug resistant (PDR), 28 were extensively drug resistant (XDR), 46 isolates were MDR, 7 were non-MDR, and 19 were pan-susceptible to all antibiotics tested. Notably, 56 isolates were non-susceptible to the 3^rd^ generation cephalosporins tested (ceftazidime and ceftriaxone), 24 were non-susceptible to carbapenems (imipenem and meropenem), and 10 were non-susceptible to the newer β-lactam/β-lactamase inhibitor, ceftazidime-avibactam.

Overall, AMR genes known to confer non-susceptibility were detected in all 100 genomes with 135 distinct alleles identified from 40 antibiotic families (**Table S1**). The majority of intrinsic *bla*_SHV_ class-A β-lactamase alleles detected were *bla*_SHV-1_ and/or *bla*_SHV-11_ (16). In 59 isolates, *bla*_SHV_ and/or *bla*_CTX-M_ ESBLs were detected, with *bla*_CTX-M-15_ (*n* = 44) being the most prevalent. Three isolates (MRSN 750999, 680172, 27106) carried *bla*_SHV-27_ as their sole ESBL gene (31), but were susceptible to the 3^rd^ generation cephalosporins. The *bla*_GES-5_ ESBL was found in a single isolate, MRSN 28183, resulting in non-susceptibility to 3^rd^ generation cephalosporins and ceftolozane-tazobactam.

Carbapenemase genes encoding IMP, KPC, NDM, OXA-48-like, and VIM enzymes were present in 24 isolates. Eleven isolates produced OXA-48-like *β*-lactamases (OXA-48, -181, -232) capable of hydrolyzing carbapenem antibiotics, with OXA-48 being the most common (*n* = 7).

All OXA-48-like positive isolates co-produced the ESBL CTX-M-15 (except a single isolate, MRSN 13748, with CTX-M-14) and as expected were non-susceptible to ceftazidime, cefepime, aztreonam, imipenem, and meropenem. Three OXA-48-like carrying isolates also co-produced NDM-1 or -5 enzymes including lineages ST-147, ST-16, and the well-studied hvKp lineage ST-23 (**Table S1**). As expected, all 10 isolates carrying genes encoding the Ambler class B Metallo-β-lactamases (MBL; IMP, NDM and VIM variants) were non-susceptible to ceftazidime-avibactam. As carbapenemase non-susceptibility can also be mediated through mutations in the outer membrane proteins (OmpK35 and/or OmpK36) in conjunction with the expression of an ESBL and/or acquired AmpC β-lactamases (32, 33), all strains were examined for known mutations in these genes. Variations in OmpK35 and/or OmpK36 were observed in 15 isolates, of which 12 carried a carbapenemase and were non-susceptible to all carbapenem antibiotics tested. The remaining three isolates had OmpK35 mutations only and were susceptible to the carbapenems. In the final panel, only 4 isolates carried an acquired AmpC β-lactamase (*bla*_FOX-5_, *bla*_DHA-1_, or *bla*_CMY-4_) and all lacked OmpK mutations. Notably, 7 isolates had a truncated *mgrB*, known to mediate colistin resistance (**Table S1**), and five were resistant (MIC > 4) to colistin by broth microdilution (BMD). As the Clinical and Laboratory Standards Institute (CLSI) guidelines (34) do not recognize susceptible breakpoints for colistin the remaining two isolates with a truncated *mgrB* (791403 and 375436) were assigned intermediate interpretation (MIC ≤0.25 and 1, respectively), as reported previously (12).

Forty-two isolates were susceptible to all three aminoglycosides tested (amikacin, gentamicin, and tobramycin) while 9 isolates were pan resistant to all aminoglycosides. All pan resistant isolates carried a 16S rRNA methyltrasferase, with the exception of MRSN 430405 for which no acquired methyltransferase gene was identified (**Table S1**). Specifically, five of the pan resistant isolates (5881, 366562, 365679, 517281, and 613682) carried methytransferase genes *rmtH, rmtF1*, or *rmtB1* and the remaining three (MRSN 27778, 607210, 368001) carried *armA*. MRSN 752729 also carried *armA* but was susceptible to amikacin and gentamicin. Reduced susceptibility was confirmed by BMD (amikacin, MIC=16; gentamicin, MIC=1). Further analysis of the *armA* sequence revealed a missense mutation at nucleotide position 617 (A to T) resulting in an amino acid substitution of isoleucine to lysine.

## Discussion

In 2017, the WHO identified ESBL and carbapenemase-resistant Enterobacteriaceae as a “Critical’ threat to human health. Similarly, the U.S. CDC named carbapenem-resistant and ESBL-producing Enterobacterales as “urgent” and “serious” threats, respectively (35). As a result, there has been a renewed and concerted effort to develop novel therapeutics and diagnostics to combat these organisms. This has been reflected at the highest level of the U.S. Government with the publication of the Presidential U.S. National Action Plan for Combating Antibiotic-Resistant Bacteria (CARB) (36). This document outlined strategies to combat this threat, including access to diverse isolates for testing. In response to these demands, the U.S. Department of Defense, through the MRSN, has published distinct panels (with corresponding metadata and genomes) for the ESKAPE pathogens *Acinetobacter baumanii* (24) and *Pseudomonas aeruginosa* (25). Herein we expand these panels by constructing a novel panel of *K. pneumoniae* isolates that, to our knowledge, is the only comprehensive panel publicly available for research and development. The panel was designed to encompass the maximum genetic diversity of the species, ensuring a diverse range of antibiotic susceptibilities, AMR genes and virulence genes.

Other panels and characterized *K. pneumoniae* strains exist, however, they mainly focus on the identification of antibiotic resistant mechanisms and were not designed to encompass the diversity of the species. For example, the U.S. CDC and FDA have collaborated to produce the AR Isolate Bank that contains multiple isolate panels for a range of bacterial pathogens and resistance mechanisms (https://wwwn.cdc.gov/arisolatebank/Overview) and this panel has proven to be an excellent resource to test the activity of antibiotic combinations (37). However, in addition to testing antibiotics, strain diversity is critical for assessing the efficacy of many emerging therapeutics including phage therapy, vaccines, and capsule polysaccharides targeted approaches (38–40). The main structural receptor for anti-*Klebsiella* phages is the external capsular polysaccharide however recent work suggests that phages also attach to other outer membrane structures below the capsule, including the O-antigen (41, 42). The panel described herein represents 54 of the 77 distinct capsule types identified by serological methods (43) and 8 predicted O serotypes (27, 44), providing a robust representation of outer membrane protein diversity to test anti-*Klebsiella* phages. Besides therapeutics, the understanding of *K. pneumoniae* pathogenesis is rapidly evolving, in particular the understanding of virulence factors that can accurately predict pathogenic potential of strains. For example, not all hvKp strains are equally virulent in murine models of infection despite carrying well-characterized virulence biomarkers (45). Herein we describe hvKp and convergent strains with diverse biomarkers to aid in these ongoing research efforts.

The epidemiology of *K. pneumoniae* over the past two decades has been characterized by widely geographically distributed “high risk” clones and the constant emergence and dissemination of new clonal groups (6). This panel not only captures the most important MDR-cKp (ST-258, ST-15, ST-11, ST-307, ST-147) and hvKp clones (ST-23, ST-380, ST-65, ST-86) currently circulating, but also encompasses the overall diversity of the species, an approach that maximizes the potential of the panel to include emerging strains, or those that may emerge in the future. To this end, the panel includes 6 novel lineages, including an XDR ST-5445 lineage carrying *bla*_CTX-M-15_, and 5 genomic convergent lineages that have not been previously described (ST-268, ST-1399, ST-48, ST-2071, ST-37). Furthermore, close attention was paid to selecting rare clones that cause localized epidemics in different regions of the world. Clones ST-43, ST-268, ST-340, ST-392 are all represented in the panel and have been reported previously as harboring NDM carbapenemases and circulating in hospitalized patients in Iran (46). Similarly, a ST-340 clone carrying a NDM carbapenemase was recovered from patients at a tertiary care hospital in South Korea (47) and infections with ST-231 and/or ST-395 clones have been identified in local hospitals in Oman (48) and South India (48, 49). A genomic surveillance study from 2013 to 2014 found ST-231, ST-340, ST-323 (carrying various ESBLs and carbapenemase genes) clones all linked to nosocomial transmission events from 4 hospitals in Melbourne Australia (50). In our panel collection the XDR clone ST-340 was collected in Asia in 2015 while the MDR clones ST-323 and ST-231 were recovered from North America in 2016 and 2018, respectively. Interestingly these localized epidemic clones have yet to globally disseminate despite being highly antibiotic resistant.

Notably, a strong association between antibiotic susceptibility and the presence of AMR genes and/or mutations was observed, with few exceptions. For example, isolates carrying *bla*_SHV-27_ ESBL had a non-ESBL phenotype. However, this discrepancy is most likely due to a base-pair substitution (A to C) in the promoter region that was previously reported in SHV-27-producing isolates susceptible to cephalosporins (51). Similarly, isolate MRSN 752729 carrying a missense mutation in *armA* 16s rRNA methyltransferase had increased susceptibility to all aminoglycoside antibiotics. Previous studies report that point mutations in *armA* can result in the inability to bind to the 16S rRNA and consequently block methylation resulting in susceptibility to aminoglycosides (52). Lastly, in this study the single GES-5-producing isolate (MRSN 28183) conferred non-susceptibility to ceftazidime, ceftriaxone, and ceftolozane-tazobactam but was susceptible to cefepime and carbapenem antibiotics. The GES-5 variant has a single amino acid substitution (G170S) compared to wild-type GES-1 and has been shown to confer activity against carbapenem antibiotics (53), yet, studies have also shown GES-5 producing *K. pneumoniae* to have minimal to no carbapenemase activity (54, 55), consistent with our observations.

In summary, this study describes the construction of a panel of 100 unique *K. pneumoniae* isolates from an extensive collection of over 3,800 *K. pneumoniae* isolates collected from across the globe. The panel encompasses the diversity of the species, includes both antibiotic susceptible and non-susceptible isolates, and captures known epidemic clones as well as sporadic ones. Furthermore, this panel captures diverse genomic convergent and hvKp strains that are rapidly emerging worldwide and are of considerable concern (15, 45). While identifying these convergent lineages does not accurately predict clinical outcomes, availability of these characterized isolates (including phylogeny, genome, and AST) will aid in the research and development of infection-control measures to improve patient care. This panel and all metadata and genomes are publicly available at no additional charge and represent an invaluable resource for genotypic and phenotypic research of this important pathogen.

## Materials and Methods

### *K. pneumoniae* repository

The MRSN collects and analyzes MDR organisms across the Military Healthcare System in the United States (23) and around the world in collaboration with the US Department of Defense’s (DoD) Global Emerging Infections Surveillance (GEIS) branch. All samples are housed in a central repository, which currently contains over 100,000 isolates, including 3,878 *K. pneumoniae* that were cultured from 2,760 patients between 2001 and 2020.

### Refinement of *K. pneumoniae* repository

To reduce redundancy in the initial 3,878 isolate set, successive isolates after the first from the same patient that shared the same ST were removed unless isolates were cultured from a different body site (e.g. urine vs blood) or were cultured >6 months apart. All isolates from the same patient with different STs were retained. This refinement resulted in a final dataset of 3,123 isolates available for analysis.

### Antibiotic susceptibility testing

AST was performed in the MRSN’s College of American Pathologists (CAP)-accredited laboratory using the Vitek 2 with the AST-95 and AST-XN09 cards (bioMerieux, NC, US). Nineteen antibiotics representing 11 different antibiotic families were tested and interpreted using Clinical and Laboratory Standards Institute (CLSI) guidelines (CLSI 2018). Susceptibility results were used to classify the isolates as pan drug resistant (PDR) (non-susceptible to all antibiotics tested), extensively drug resistant (XDR) (non-susceptible to ≥1 agent in all but ≤2 families), MDR (non-susceptible to ≥1 agent in ≥3 antibiotic families), and non-MDR (non-susceptible to 1 or 2 categories) using a modification of the criteria defined by Magiorakos et al (30). When necessary, MICs were repeated in triplicate using broth microdilution and CLSI guidelines (CLSI 2018).

### Whole-genome sequencing and data analysis

Briefly, isolates were sequenced on an Illumina MiSeq or NextSeq benchtop sequencer (Illumina, Inc., CA, US) and analyzed as previously described (24). Where appropriate, long read sequencing was performed with the Oxford nanopore MinION sequencer (Oxford Nanopore Technologies), as previously described (12). *In silico* MLST typing, virulence loci, polysaccharide capsule (K) loci, and lipopolysaccharide (O) loci typing were performed using Kleborate v2.0.1 (56). Novel MLST STs were determined using the Klebsiella PasteurMLST sequence database (https://bigsdb.pasteur.fr/klebsiella).

AMRFinderPlus v3.9.8 (57) and ARIBA v2.14.4 (58) were used to identify resistance alleles. Basic assembly statistics are available (see **Table S2** in the supplemental material).

### cgMLST analysis

The draft genomes of all 3,878 *K. pneumoniae* isolates were uploaded and analyzed using Ridom SeqSphere+ (59) using the *K. pneumoniae* cgMLST scheme (https://www.cgmlst.org/ncs). To be included in the analysis, isolates had to contain 90% of the 2,358 genes included in the cgMLST scheme. The resulting minimum spanning tree (MST) was then used to select 346 strains that capture the diversity of the strain collection.

### Core-genome SNP analysis

PanSeq (60) was run with a fragmentation size of 500 bp to find sequences with ≥95% identity in ≥95% of the isolates to generate the core genome single nucleotide polymorphism (SNP) alignment for the initial set of 346 isolates. RAxML (v8.2.11) (61) was used to generate a phylogenetic tree for the core SNP alignment. The SNP-based phylogeny was built from a 317-kb variable position alignment using the general time reversible (GTR) GAMMA model and the rapid bootstrapping option for nucleotide sequences, using 100 replicates. Using this approach, 100 strains were selected to represent the final diversity panel. For the final diversity set of 100 isolates, reads were checked for contamination at the species level with Kraken2 (v2.0.8-beta) (62) and at the strain level using ConFindr (v0.4.8) (63) with parameters bf=0.05 and q=30, as previously described (24). A phylogenetic tree of the 100 isolates was constructed with PanSeq and RAxML as described above. The SNP-based phylogeny was built from 169-kb variable position alignment. For all 100 isolates included in the panel, genome annotations were performed using NCBI Prokaryotic Genome Annotation Pipeline (v4.8) and core and pangenomes were calculated with Roary (v3.12.0) (64). The final 100 genomes have been deposited in the National Center for Biotechnology Information under BioProject PRJNA717739.

### Diversity panel availability

The final *K. pneumoniae* diversity panel has been deposited at BEI resources (https://www.beiresources.org/) and is currently available for research purposes under catalogue #NR-55604.

## Acknowledgements

This study was funded by the U.S. Army Medical command, the Defense Medical Research and Development Program and the Department of Defense (DoD) Global Emerging Infections Surveillance (GEIS) program (Award P0140_20_WR_04 to J.W. Bennett and P.T. McGann).

We gratefully acknowledge the participation of clinical laboratories across the Military Healthcare System, GEIS-affiliated overseas laboratories, and numerous collaborators for their contributions of bacteria to the MRSN repository.

The manuscript has been reviewed by the Walter Reed Army Institute of Research and there is no objection to its presentation. The opinions or assertions contained herein are the private views of the authors and are not to be construed as official or reflecting the views of the Department of the Army or the Department of Defense.

